# ACE2 and SARS-CoV-2 Expression in the Normal and COVID-19 Pancreas

**DOI:** 10.1101/2020.08.31.270736

**Authors:** Irina Kusmartseva, Wenting Wu, Farooq Syed, Verena Van Der Heide, Marda Jorgensen, Paul Joseph, Xiaohan Tang, Eduardo Candelario-Jalil, Changjun Yang, Harry Nick, Jack L. Harbert, Amanda Posgai, Richard Lloyd, Sirlene Cechin, Alberto Pugliese, Martha Campbell-Thompson, Richard S. Vander Heide, Carmella Evans-Molina, Dirk Homann, Mark A. Atkinson

## Abstract

Diabetes is associated with increased mortality from Severe Acute Respiratory Syndrome Coronavirus-2 (SARS-CoV-2). Given literature suggesting a potential association between SARS-CoV-2 infection and diabetes induction, we examined pancreatic expression of the key molecule for SARS-CoV-2 infection of cells, angiotensin-converting enzyme-2 (ACE2). Specifically, we analyzed five public scRNAseq pancreas datasets and performed fluorescence *in situ* hybridization, Western blotting, and immunolocalization for ACE2 with extensive reagent validation on normal human pancreatic tissues across the lifespan, as well as those from coronavirus disease 2019 (COVID-19) patients. These *in silico* and *ex vivo* analyses demonstrated pancreatic expression of ACE2 is prominent in pancreatic ductal epithelium and the microvasculature, with rare endocrine cell expression of this molecule. Pancreata from COVID-19 patients demonstrated multiple thrombotic lesions with SARS-CoV-2 nucleocapsid protein expression primarily limited to ducts. SARS-CoV-2 infection of pancreatic endocrine cells, via ACE2, appears an unlikely central pathogenic feature of COVID-19 as it relates to diabetes.

## INTRODUCTION

The coronavirus disease 2019 (COVID-19) pandemic caused by Severe Acute Respiratory Syndrome-Coronavirus-2 (SARS-CoV-2) has created a global healthcare crisis (Mercatelli and Giorgi, 2020). With infections continuing to rise in many countries and the potential for continuing viral persistence in the absence of a vaccine, there is an urgent need to better understand SARS-CoV-2-mediated pathology. Key to this are efforts examining human tissues potentially susceptible to infection.

While initial reports primarily focused on pulmonary and cardiovascular manifestations, other organs including the kidney, brain and intestines, as well as the pancreas, have since been noted as affected by this disorder’s pathophysiology (Connors and Levy, 2020; Fox et al., 2020; Hanley et al., 2020; Lee et al., 2020b; Liu et al., 2020; Menter et al., 2020; Rapkiewicz et al., 2020; Varga et al., 2020; Wang et al., 2020; Wichmann et al., 2020). Indeed, recent reports have raised the question of whether SARS-CoV-2 might infect the pancreas and possibly potentiate or exacerbate diabetes in either of its predominant forms, type 1 or type 2 diabetes (i.e., T1D or T2D, respectively). These studies noted elevated serum levels of the exocrine pancreatic enzymes, amylase and lipase, as well as development or worsening of hyperglycemia in SARS-CoV-2 positive individuals (Wang et al., 2020), high prevalence of diabetic ketoacidosis in hospitalized COVID-19 patients (Goldman et al., 2020; Li et al., 2020), increased COVID-19 mortality in patients with T1D and T2D (Barron et al., 2020; Holman et al., 2020), increased incidence of new-onset T1D in specific geographic clusters (Unsworth et al., 2020), case reports linking the timing of T1D onset to COVID-19 (Marchand et al., 2020), and pancreatic expression of angiotensin-converting enzyme-2 (ACE2), through which SARS-CoV-2 gains access to cells (Chen and Hao, 2020), potentially including in insulin-producing β-cells (Lee et al., 2020b; Yang et al., 2020). These reports have collectively led to the hypothesis that SARS-CoV-2 expression in β-cells may potentiate or exacerbate T1D or T2D. However, ACE2 expression in the human pancreas is only partially characterized, with conflicting results in both the endocrine and exocrine compartments (Fignani et al., 2020; Hikmet et al., 2020; Lee et al., 2020b; Yang et al., 2010; Yang et al., 2020). Indeed, the most cited report (Yang et al., 2010) utilized a single reagent (e.g., anti-ACE2 antibody), was limited in number of tissue samples/cases, and lacked reagent validation.

To better understand the potential impact of SARS-CoV-2 on diabetes, we carried out an extensive investigation of the human pancreas, with particular focus on its endocrine component, the islet of Langerhans. Specifically, we performed an integration-analysis of publicly available single cell RNA sequencing (scRNAseq) data from isolated human islets, and coupled these findings with direct visualization of gene and protein expression for ACE2 using single molecular fluorescence *in situ* hybridization (smFISH), chromogen-based immunohistochemistry (IHC), and multicolor immunofluorescence (IF) in human tissue. Importantly, we employed four commercially available ACE2 antibodies and included validation studies by IHC and immunoblot using known ACE2 positive tissues. Finally, we analyzed SARS-CoV-2 nucleocapsid protein (NP) expression in autopsy-derived tissues from deceased COVID-19 patients to assess whether the virus was detected in pancreatic islet endocrine cells.

## RESULTS AND DISCUSSION

### Low *ACE2* and *TMPRSS2* Gene Expression in Human Pancreatic Endocrine Cells

Diabetes, obesity, and advanced age increase the risk of COVID-19 mortality (Zhou et al., 2020). Autopsy studies of SARS-CoV-2 infected individuals demonstrate systemic viral dissemination with persistence in multiple organs including lungs and kidneys (Hanley et al., 2020; Liu et al., 2020; Menter et al., 2020; Wichmann et al., 2020). Autopsy studies of pancreas have been limited, likely due to challenges related to its post-mortem autolysis and presumed limited clinical significance for COVID-19. However, recent studies (Barron et al., 2020; Fignani et al., 2020; Goldman et al., 2020; Holman et al., 2020; Li et al., 2020; Marchand et al., 2020; Unsworth et al., 2020; Wang et al., 2020) spurred interest in ACE2 expression in the pancreas, particularly the endocrine compartment, to address a potential relationship between diabetes and COVID-19.

SARS-CoV-2 entry into cells via ACE2 can be facilitated by the mucosal serine proteases, TMPRSS2 and TMPRSS4 (Lee et al., 2020b; Zang et al., 2020). Therefore, we investigated expression patterns of these molecules in isolated human islets by conducting an integrated analysis of scRNAseq data from five datasets including 22 non-diabetic and 8 T2D individuals (Baron et al., 2016; Grün et al., 2016; Lawlor et al., 2017; Muraro et al., 2016; Segerstolpe et al., 2016). This analysis revealed low frequency of *ACE2* expressing cells and low *ACE2* expression levels in the majority of islet cell subsets (**Fig. 1A,B**). In non-diabetic donors, *ACE2* was expressed in <2% of endocrine, endothelial, and immune cells. *ACE2* was detectable in 4.11% of acinar cells and 5.54% of ductal cells in non-diabetic donors and 8.07% of acinar and 8.13% of ductal cells in donors with T2D (**Table S1**). Expression levels of *ACE2* were not different between non-diabetic donors and donors with T2D in any of the islet cell subtypes. *TMPRSS2* was detectable in 53.73% of acinar and 50.55% of ductal cells in non-diabetic donors, and 71.43% of acinar and 58.74% of ductal cells in donors with T2D (**Table S1**). Apart from α-cells, which demonstrated 16.55% positivity in non-diabetic donors, *TMPRSS2* expression was low in the majority of endocrine cell subsets (**Fig. 1C**). *TMPRSS2* showed higher relative expression levels in ductal and acinar cells compared to β-cells (adjusted *P* = 1.31×10^−291^ and *P* < 1×10^−300^, respectively), and was detectable in an elevated proportion of these cells in non-diabetic donors (*P* < 0.05) (**Fig. 1D**). Neither *ACE2* nor *TMPRSS2* expression differed significantly in β-cells from non-diabetic versus T2D donors (**Fig. S1A,B**).

**Fig. 1.**
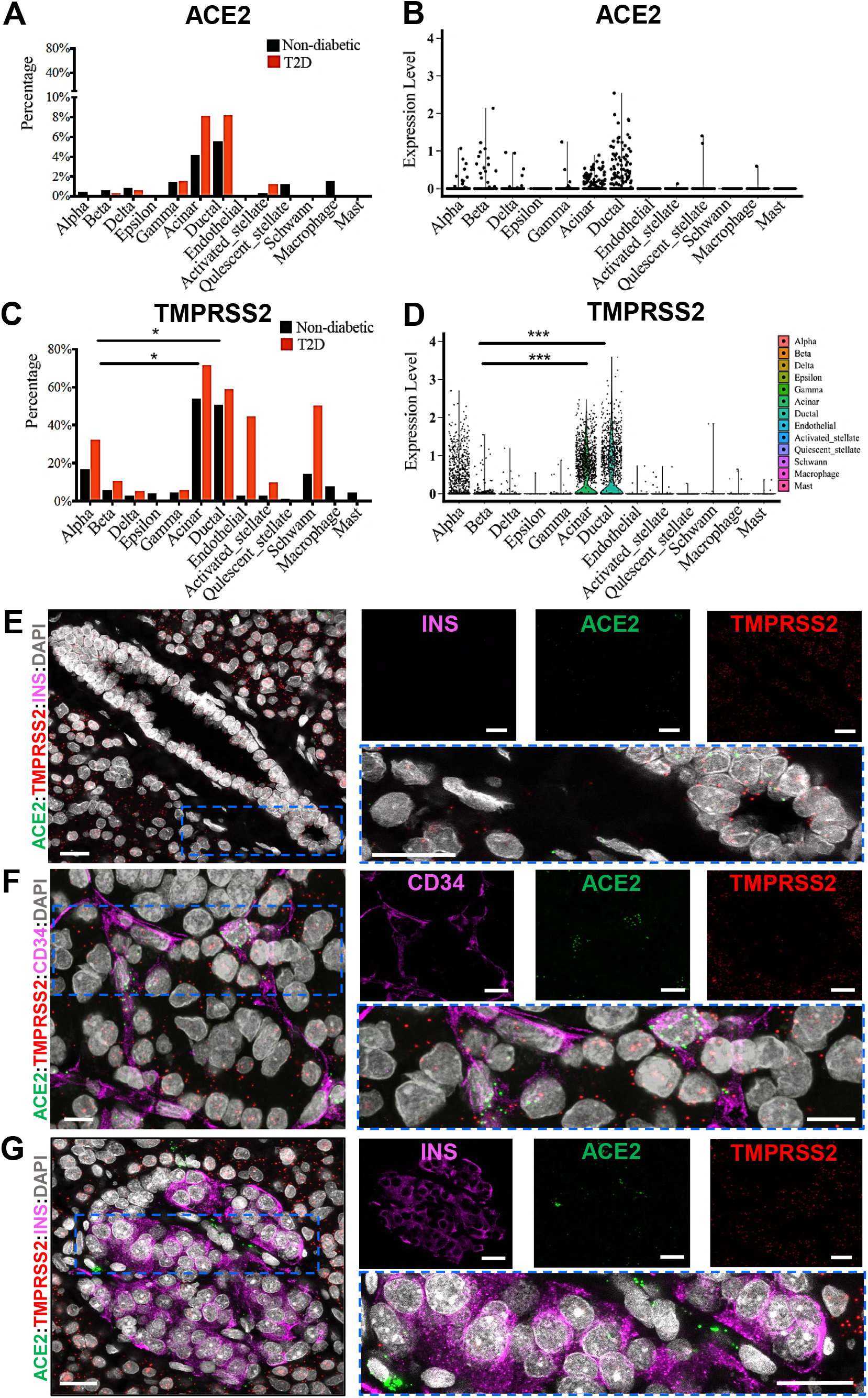
SARS-CoV-2 associated gene expression in human pancreas. (A) Bar graph showing the percentage of cells with detectable *ACE2* in islets from pancreata of donors with (n=2,705 cells) and without type 2 diabetes (n=12,185 cells). (B) Violin plot showing the distribution of *ACE2* normalized expression in islet cells from pancreata of non-diabetic donors. (C) Bar graph showing the percentage of cells with detectable *TMPRSS2* in islets isolated from pancreata of donors with (n=2,705 cells) and without type 2 diabetes (n=12,185 cells); \**P*<0.05, paired t-test for indicated comparisons. (D) Violin plot showing the distribution of *TMPRSS2* normalized expression in islets cells from pancreata of non-diabetic donors; ***adjusted *P*<0.001, Wilcoxon rank sum tests. (E) Representative images of smFISH for *ACE2* and *TMPRSS2* mRNA in human pancreatic tissue sections counter stained for insulin (magenta). Inset highlights mRNA distribution in pancreatic ducts; scale bar: 20µm. (F) Representative smFISH images showing the presence of *ACE2* and *TMPRSS2* in CD34 positive cells in human pancreatic tissue sections; scale bar: 10µm. (G) Representative images of smFISH for *ACE2* and *TMPRSS2* mRNA in human pancreatic tissue sections counter stained for insulin (magenta). Inset highlights distribution in the endocrine pancreas; scale bar: 20µm. See also **Table S1, Fig. S1, Table S2**, and **Fig. S2A**.

We also investigated expression patterns for other SARS-CoV-2 associated genes, including *TMPRSS4, TMPRSS11D, CTSL* and *ADAM17* (**Table S1** and **Fig. S1C-H**). Similar to *TMPRSS2, TMPRSS4* expression was enriched in acinar and ductal cells. While *TMPRSS4* was expressed in a similar proportion of α- and β-cells, relative expression levels tended to be low in the endocrine pancreas (**Fig. S1C-D**). *CTSL* and *ADAM17* were detected at higher levels in α- and β-cells (**Fig. S1E-F**), while *TMPRSS11D* expression was low in most cell types examined (**Fig. S1G**). Within the β-cells, only *CTSL* showed higher expression in donors with T2D compared to donors without diabetes (adjusted *P* = 8.94X 10-32, **Fig. S1H**).

To directly visualize *TMPRSS2* or *ACE2* mRNA expression patterns, we used smFISH to analyze non-diabetic, SARS-CoV-2 negative, “normal” human pancreata from six donors across a wide age-span (**Table S2** and **Fig. 1E-G**) with duodenum, ileum, and kidney used as positive controls (**Fig. S2A**). Results from smFISH were consistent with scRNAseq analyses. Specifically, *ACE2* mRNA was observed at low frequency in the pancreas, its signal mostly localized to acinar, ductal and CD34+ endothelial cells (**Fig. 1E-F**). *TMPRSS2* showed a similar pattern but was expressed at higher frequency. Finally, in the islets, we observed limited expression of *TMPRSS2* or *ACE2* in insulin positive β-cells (**Fig. 1G**).

### Extensive Reagent Validation to Detect ACE2 Protein Expression

Information regarding ACE2 protein expression in pancreatic tissue sections remains limited and unfortunately contradictory. A 2010 study on SARS-CoV and its relationship with diabetes interrogated ACE2 expression in a single donor (43 years of age), with an unspecified antibody, and reported weak ACE2 staining in exocrine tissues but pronounced expression in pancreatic islets (Yang et al., 2010). A recent preprint (Fignani et al., 2020) described heterogenous ACE2 expression across donors, pancreatic lobes and islets, and identified three main ACE2-expressing cell types in formalin fixed, paraffin embedded (FFPE) pancreatic sections from seven donors (aged 22-59 years) probed with a single ACE2 antibody (MAB933): endothelial cells/pericytes, ductal cells, and in an analysis of 128 islets, β-cells that often presented with a granular staining pattern, partially overlapping with insulin. In contrast, an analysis of tissue microarrays containing pancreatic FFPE sections from 10 donors (aged 30-79 years) utilizing two ACE2 antibodies (MAB933 and HPA000288) reported ACE2 expression restricted to endothelial cells/pericytes and interlobular ducts while ACE2 was not detectable in islets, acinar glandular cells, intercalated ducts, or intralobular ducts (Hikmet et al., 2020). This study, together with a preprint employing six ACE2 antibodies (Lee et al., 2020a), is noteworthy for the broad range of major human tissues and organs investigated as well as the delineation of mostly shared but also, some distinctive ACE2 antibody staining properties. In light of these diverging observations, it is imperative to utilize multiple ACE2 antibodies on larger donor cohorts to gather a comprehensive description of ACE2 expression in the pancreas.

We selected four widely referenced, commercially available antibodies recognizing specific epitopes of ACE2 (**Fig. 2A**), to evaluate by immunoblot (**Fig. 2B** and **Fig. S2B-C**) using protein extracts from three non-diabetic, SARS-CoV-2 negative, “normal” pancreas donors (**Table S2**). ACE2 is an 805 amino acid protein (UniProt Q9BYF1) with a theoretical molecular mass of 94.2 kDa but an actual mass of ∼120 kDa due to glycosylation at N-terminus sites (Tipnis et al., 2000). Accordingly, a pronounced ∼120kDa band was readily visualized by AF933 and ab108252 while ab15348 and especially MAB933 revealed a much weaker band (**Fig. 2B** and **Fig. S2B**). In our hands, all four antibodies demonstrated robust and essentially commensurate staining in human duodenum and kidney FFPE sections (**Fig. S2D**), consistent with (Hikmet et al., 2020; Lee et al., 2020a).

**Fig. 2.**
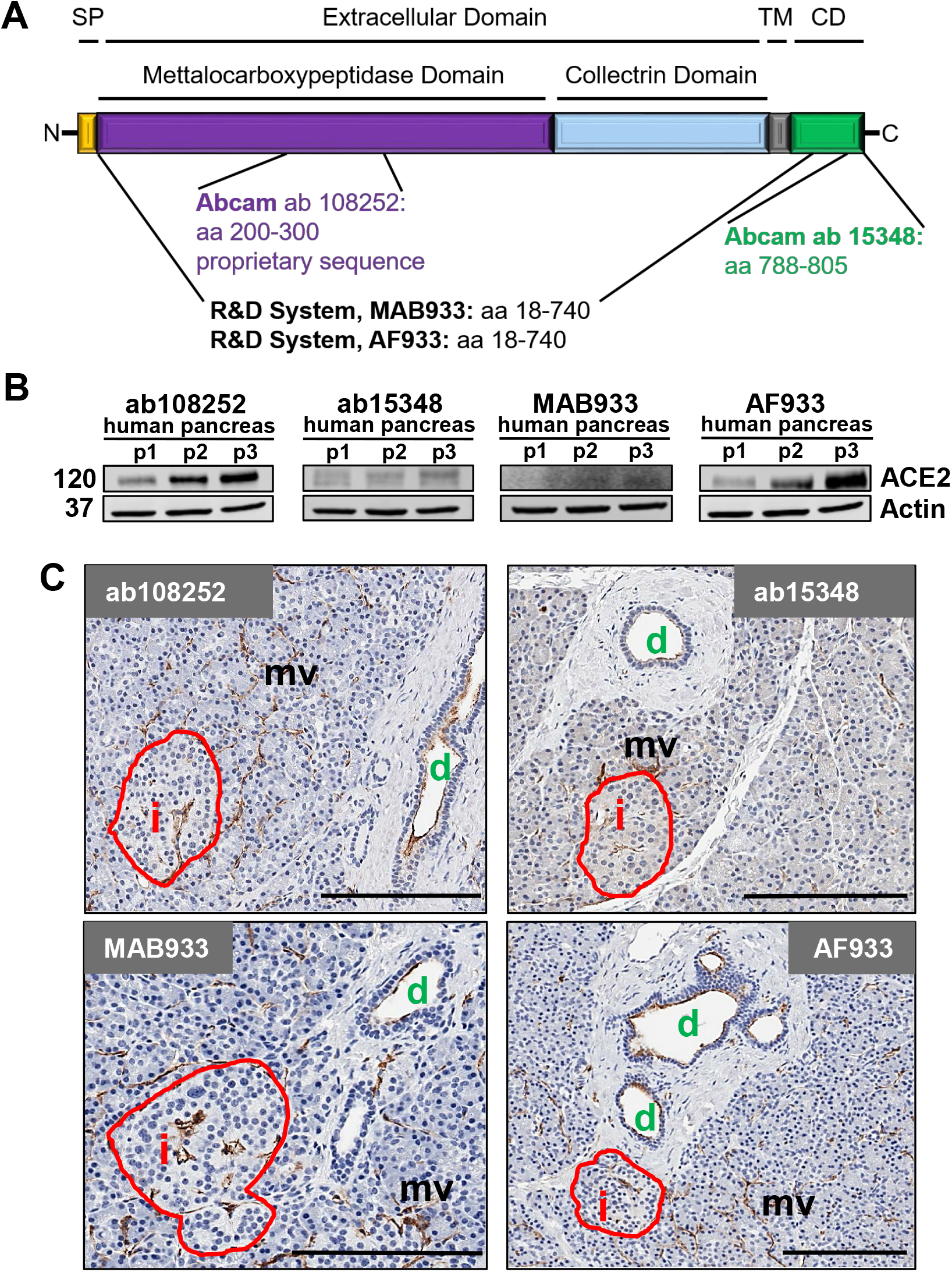
ACE2 protein is readily detected in normal human pancreas with its expression largely restricted to ductal and microvascular structures. (A) ACE2 protein structure illustrating the location of respective antibody directed antigen sites: SP, signal peptide; TM, transmembrane domain; CD, cytoplasmic domain. (B) Immunoblot analysis of four commercially available ACE2 antibodies using total pancreas lysates from three control organ donors (p1-p3) with accompanying Actin labeling. (C) Representative IHC images of human pancreas tissue sections stained for ACE2 using four commercially available ACE2 antibodies. Scale bars: 200 µm. Abbreviations: i, islet; d, duct; mv, microvasculature. See also **Table S2** and **Fig. S2B-D**.

We next visualized ACE2 via chromogen-based IHC in FFPE pancreata from non-diabetic, SARS-CoV-2 negative “normal” donors using all four antibodies and consistently observed positive staining in the microvasculature and ductal epithelium (**Fig. 2C**). To further validate specificity, we utilized a peptide-blocking assay wherein ab108252 was pre-incubated with an ACE2 peptide prior to IHC staining (**Fig. S2E**). Taking Western blot and IHC validation into account, ab108252 produced a clearly detectable 120kDa band and crisp *in situ* staining that was completely blocked by pre-incubation with the ACE2 peptide. This is a monoclonal antibody, which offers a high degree of specificity and consistency between lots. Hence, we elected to use ab108252 in subsequent IHC and IF assays.

### Human Pancreatic Expression of ACE2 Protein is Primarily Localized to Duct Epithelium and Microvasculature Across the Lifespan

To quantitatively evaluate pancreatic ACE2 protein expression, a FFPE tissue cross-section from each of 36 SARS-CoV-2 negative donors without diabetes (aged 0-72 years, **Table S2**) was stained for insulin and ACE2 (ab108252) and scanned to produce a whole-slide image. Co-localization of ACE2 with insulin was not observed (**Fig. S3A-C**). The tissue area staining positive for ACE2 was analyzed using the HALO Area Quantification algorithm (**Fig. 3A**). Donors were binned into six age groups: neonate (0-0.25 years, n=6), infant/toddler (0.25-2 years, n=6), child (2-11 years, n=6), adolescent (11-15 years, n=6), young adult (20-35 years, n=6), and senior adult (51-72 years, n=6). Collectively, these data demonstrate the percentage of tissue staining positive for ACE2 increases steadily from birth throughout childhood, peaking in adolescence and maintained through early adulthood, followed by a decline in persons over 50 years of age (**Fig. 3B**).

**Fig. 3.**
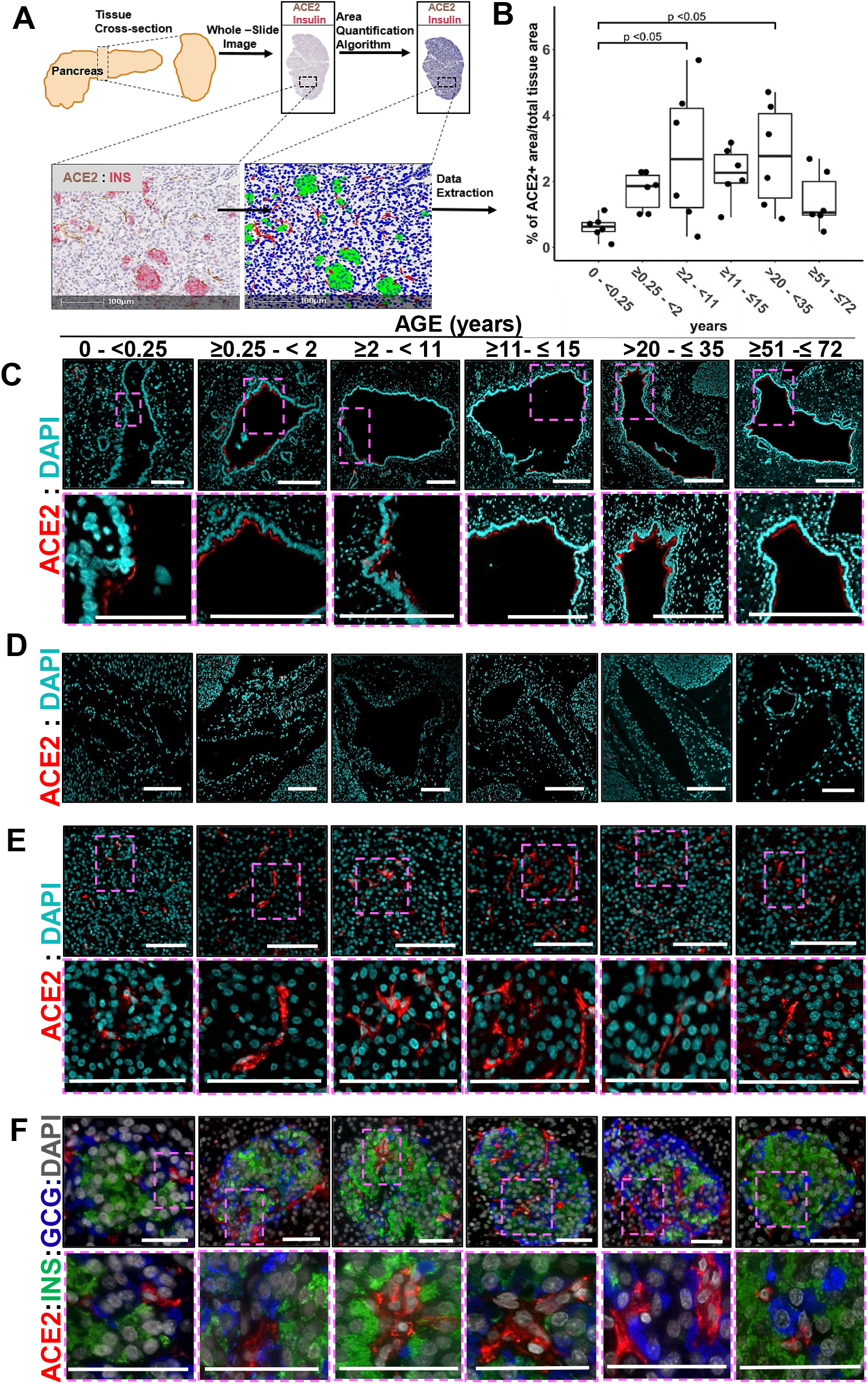
ACE2 protein expression is evident in normal pancreata throughout the human lifespan. (A) Scheme of the experimental setup illustrating human pancreatic tissue processing, whole stained slide imaging, and machine learning algorithm application for the generation of ACE2 protein expression data. (B) Quantification of ACE2 protein expression in the pancreas of control organ donors with ages ranging from birth to 72 years shows progressive developmental changes. Data are presented as mean±SD with analysis by one-way ANOVA and Tukey’s post hoc test for multiple comparisons. (C) Representative confocal images of ACE2 protein expression in pancreatic ducts of control donors across different age groups. Scale bars (left to right): 100µm, 200µm, 200µm, 300µm, 300µm, 300µm. (D) Representative immunofluorescence images showing lack of ACE2 protein expression in pancreatic blood vessels from control donors across different age groups. Scale bars (left to right): 100µm, 200µm, 200µm, 200µm, 200µm, 100µm. (E) Representative immunofluorescence images show ACE2 protein expression in pancreatic microvasculature from control donors across different age groups. Scale bars: 100µm. (F) Representative immunofluorescence images of pancreatic islets showing ACE2 protein expression restricted to the islet’s microvasculature in pancreata from control organ donors across different age groups. Scale bars: 50µm. See also **Table S2** and **Fig. S3**.

To more precisely visualize pancreatic ACE2 localization, we performed IF staining for ACE2 (again using ab108252) in conjunction with insulin and glucagon (**Fig. 3C-F**) or with CD34 (**Fig. S3D-F)**. Across all ages, we observed ACE2 expression in the pancreatic ductal epithelium (**Fig. 3C**) but not major blood vessels (**Fig. 3D**), based on their morphology and geographic positioning. ACE2 was highly expressed in microvasculature within acinar and islet regions, with no evidence of α-cells or β-cells expressing ACE2 (**Fig. 3E,F** and **Fig. S3**). Our data corroborate findings by one group (Hikmet et al., 2020), yet contrast with two others (Fignani et al., 2020; Yang et al., 2010). These disparate results may be due to technical (e.g., antigen retrieval, reagent), material (e.g., isolated islets versus tissue sections), or donor differences. Though one cannot definitively exclude the possibility for ACE2 protein expression in endocrine islet cells, our analysis of 36 donors across a wide age range provides a comprehensive view of pancreatic ACE2 localization and is independently corroborated by scRNAseq and smFISH gene expression data (**Fig. 1**).

### SARS-CoV-2 NP Localized to Pancreatic Ductal Epithelium from COVID-19 Patients With and Without T2D

Following autopsy, pancreatic pathology was reviewed by hematoxylin and eosin (H&E) stained sections in three patients with fatal COVID-19 (aged 45-72 years, **Fig. 4A-C**), two of whom had a previous diagnosis of T2D (**Table S3**). In Patient 1, who did not have diabetes, major findings included severe fatty replacement of acinar cell mass and moderate arteriosclerosis (**Fig. 4A**). Lobules contained fibrotic centers with residual acinar cells and islets surrounding ductules (**Fig. 4A insert)**. Islets were observed primarily within fibrotic regions. Patient 2 had moderate fatty replacement and limited centrolobular fibrosis (**Fig. 4B**). Dystrophic calcification of adipocytes was rare. Numerous islets were observed. One microthrombus was observed without adjacent hemorrhages (**Fig. 4B insert**). Patient 3 showed mild to moderate arteriosclerosis with acinar regions containing mild centrolobular fibrosis (**Fig. 4C**). Moderate numbers of islets were present of varying sizes (**Fig. 4C insert**). None of the patients showed islet amyloidosis or acute polymorphonuclear cell infiltrates. These histopathological findings were compatible with the normal range of expected lesions within the exocrine compartment in pancreata from aged patients and those with T2D.

**Fig. 4.**
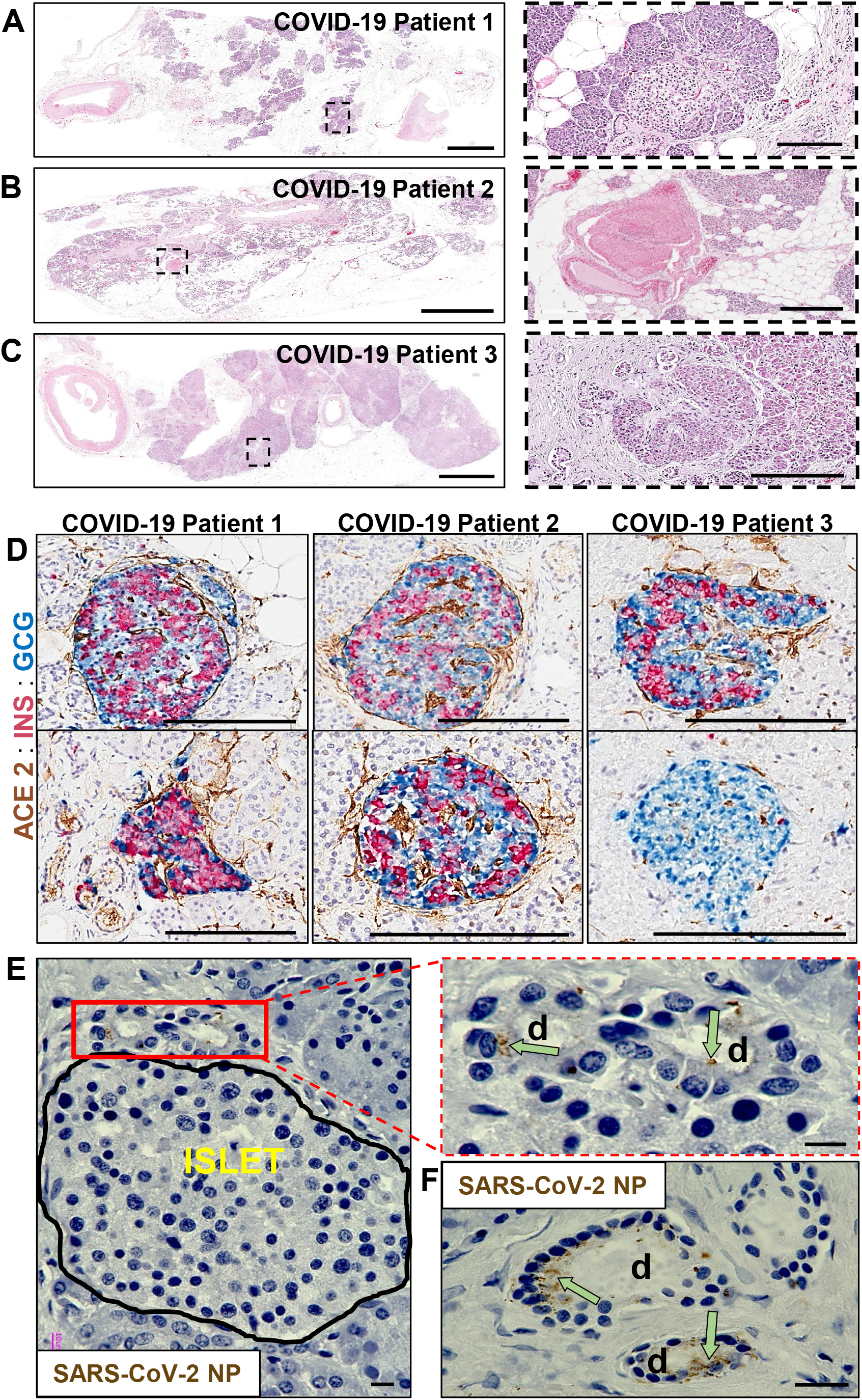
Pathological changes in pancreata of COVID-19 patients. (A) Pancreas tissue section from COVID-19 Patient 1 stained for H&E. Inset highlights fibrotic center with residual acinar cells and islet surrounding ductules. Scale bars: 3mm, inset 200µm. (B) Pancreas tissue section from COVID-19 Patient 2 stained for H&E. Inset highlights microthrombus without adjacent hemorrhages. Scale bars: 4mm, inset 400µm. (C) Pancreas tissue section of COVID-19 Patient 3 stained for H&E. Inset highlights a large, irregularly shaped pancreatic islet surrounded by fibrotic tissue. Scale bars: 4 mm, inset 200µm. (D) Representative pancreas tissue sections from three COVID-19 patients stained for ACE2, insulin (INS) and glucagon (GCG). Scale bars: 200µm. (E) SARS-CoV-2 NP observed in intralobular ducts (d) near an islet in the pancreas of COVID-19 Patient 1. Scale bars: 10µm. (F) Representative image of multiple ducts showing SARS-CoV-2 NP positivity in the pancreas of COVID-19 Patient 1. Scale bar: 20µm. See also **Table S3** and **Fig. S4**.

IHC showed islets containing insulin-positive (INS+) β-cells with mostly spherical profiles, small to medium sizes, and varying proportions of β- to α-cells in all three patients (**Fig. 4D**). Numerous single and clustered INS+ and glucagon-positive (GCG+) cells were observed in ductules of fibrotic foci. The endothelium showed moderate ACE2 staining intensity within both endocrine and exocrine compartments. The ductal epithelium showed low to moderate ACE2 staining intensity throughout the cytoplasm similar to that observed in non-diabetic pancreas donors (**Fig. 3**).

IHC for SARS-CoV-2 NP was also conducted to investigate the cellular distribution of the virus. A lung sample from a patient with COVID-19 pneumonia was used to optimize staining conditions. Immunopositive alveolar epithelial cells and macrophages were observed with numerous viral inclusions (**Fig. S4A-B**). In Patient 1 SARS-CoV-2 NP was present in some intralobular and interlobular ductal epithelial cells shown near an islet and widely scattered throughout the exocrine regions (**Fig. 4E,F** and **Fig. S4C**). Pancreata from Patients 2 and 3, who had T2D, showed little to no immunopositivity for SARS-CoV-2 NP.

We believe these data provide an important foundation for considerations of pancreatic SARS-CoV-2 infection as a potential trigger for diabetes. However, the histopathology data presented herein do not support a causative link between the two conditions via ACE2-mediated *in vivo* infection of β-cells with SARS-CoV-2. Preferential ACE2 gene and protein expression in microvascular and ductal structures suggest these cells may constitute a more likely target for viral infection of islets rather than endocrine cells. Indeed, based on our observations from three COVID-19 individuals, direct SARS-CoV-2 infection of the pancreas occurred to a very limited degree within pancreatic ductal epithelium but not islet cells, and was not associated with polymorphonuclear infiltrations.

### Limitations of Study

In theoretical conflict with our efforts, Yang *et al*. reported ACE2 protein expression in endocrine cells of isolated human islets and demonstrated their susceptibility to infection with SARS-CoV-2 (Yang et al., 2020). With both public scRNAseq data and our *in situ* smFISH experiments documenting the presence of *ACE2* mRNA in small subsets of pancreatic endocrine cells, it remains unclear whether this forms an extremely limited basis for susceptibility to SARS-CoV-2 infection. However, it remains unknown whether the process of islet isolation may influence endocrine cell ACE2 expression or whether viral dosage might influence their ability to undergo SARS-CoV-2 infection *ex vivo*. Contrasting epidemiological reports from the United Kingdom and Germany (Tittel et al., 2020; Unsworth et al., 2020) not only underscore the requirement for data on diabetes incidence and SARS-CoV-2 infection rates in defined populations over time, but also raise the need for studying pancreatic tissues from a variety of geographic populations.

## SUPPLEMENTAL INFORMATION

Supplemental Information includes 4 Figures and 3 Tables.

## AUTHOR CONTRIBUTIONS

IK researched data, generated figures, and wrote the manuscript; WW, FS, VvdH, MJ, PJ, XT, ECJ, and CY researched data and reviewed/edited the manuscript; HN generated figure 2A, contributed to discussion, and reviewed/edited the manuscript, JLH reviewed pathology and reviewed/edited the manuscript, ALP contributed to discussion and wrote the manuscript; RL, SC, and AP contributed to discussion and reviewed/edited the manuscript, RSVH procured COVID-19 autopsy tissues, reviewed pathology, and reviewed/edited the manuscript, MCT reviewed pathology and wrote the manuscript, CEM, DH, and MAA conceived of the study and wrote the manuscript.

## ACKNOWLEDGEMENTS

We thank the families of the organ donors and autopsy patients for the gift of tissues.

## STAR METHODS

### RESOURCE AVAILABILITY

#### Lead Contact

Further information and requests for reagents may be directed to and will be fulfilled by the lead contact/corresponding author, Mark A Atkinson.

#### Materials Availability

This study did not generate new unique reagents. Tissues used in this study were obtained from the Network for Pancreatic Organ donors with Diabetes (nPOD) and from autopsies performed on deceased COVID-19 patients. nPOD tissues are freely available to approved investigators following successful application to the nPOD Tissue Prioritization Committee (TPC).

#### Data and Code Availability

This study did not generate code. Single cell sequencing data were obtained from the Gene Expression Omnibus (GEO) Repository: GSE84133 (Baron et al., 2016), GSE81076 (Grün et al., 2016), GSE85241 (Muraro et al., 2016), and GSE86469 (Lawlor et al., 2017), as well as from ArrayExpress: accession number E-MTAB-5061 (Segerstolpe et al., 2016). Original histology and additional de-identified organ donor data are available from nPOD at the nPOD online digital pathology database and from the corresponding author upon reasonable request.

## EXPERIMENTAL MODEL AND SUBJECT DETAILS

### nPOD Donors and Sample Processing

Transplant-quality pancreas, duodenum, and kidney were recovered by JDRF nPOD (www.jdrfnpod.com) from 36 COVID-19 negative organ donors without diabetes (**Table S2**) according to established protocols and procedures (Campbell-Thompson et al., 2012), as approved by the University of Florida Institutional Review Board (201400486), the United Network for Organ Sharing (UNOS), and according to federal guidelines with informed consent obtained from each donor’s legal representative. Organs were shipped in transport media on ice via organ courier to the nPOD Organ Pathology and Processing Core (OPPC) at the University of Florida where tissues were processed (Campbell-Thompson et al., 2012). Medical chart and medical-social questionnaire reviews were performed, and T1D-associated autoantibodies measured by ELISA (Wasserfall et al., 2016) to confirm non-diabetic health status. Donor demographics, hospitalization duration, and organ transport time were determined from hospital records or UNOS.

### Autopsy Subjects and Sample Processing

Pancreas was recovered from three patients who tested positive for SARS-CoV-2 by reverse transcription polymerase chain reaction (RT-PCR) test within 24-48 hours of death at the University Medical Center New Orleans (New Orleans, LA), which is equipped with an autopsy suite that meets U.S. Centers for Disease Control and Prevention standards for autopsy of patients with COVID-19 (**Table S3**). Consent for autopsy without restriction was given by each patient’s next of kin, and the studies within this report were determined to be exempt from oversight by the Institutional Review Board at Louisiana State University Health Sciences Center.

## METHOD DETAIL

### Single Cell RNA-sequencing (scRNAseq) Data Analysis

Five human islet scRNA-seq datasets were obtained from publicly available repositories. These included four datasets from the Gene Expression Omnibus (GEO) Repository: GSE84133 (inDrop) (Baron et al., 2016), GSE81076 (Celseq) (Grün et al., 2016), GSE85241 (CelSeq2) (Muraro et al., 2016), and GSE86469 (Fluidigm C1) (Lawlor et al., 2017). In addition, we analyzed an ArrayExpress database under the accession number E-MTAB-5061 (SMART-Seq2) (Segerstolpe et al., 2016). For all scRNAseq datasets, the same initial normalization was performed: gene expression values for each cell were divided by the total number of transcripts and multiplied by 10,000. Following log-transformation, cells were filtered that expressed fewer than 500 genes/cell (InDrops), 1,750 genes/cell (CelSeq), or 2,500 genes/cell (CelSeq2, Fluidigm C1, and SMART-Seq2) in accordance with the methods employed in the original corresponding publications, leaving 14,890 cells in total for the combined analysis. Pancreatic islet cell subtypes were identified using methods outlined in (Butler et al., 2018).

To integrate scRNAseq data, we applied canonical correlation analysis (CCA) in Seurat v.3 (Butler et al., 2018) using “FindIntegrationAnchors” and “IntegrateData” functions. We chose the top 2,000 variable genes from each dataset to calculate the correlation components (CCs) and “FindClusters” was utilized for shared nearest neighbor (SNN) graph-based clustering. Clusters were visualized with t-distributed stochastic neighbor embedding (t-SNE) by running dimensionality reduction with “RunTSNE” and “TSNEPlot”. To compare the average gene expression within the same cluster between cells of different samples, we applied the AverageExpression function. Statistical analyses are further described in QUANTIFICATION AND STATISTICAL ANALYSIS below. Violin plots (VlnPlot) were used to visualize gene expression levels (**Fig. 1A-D** and **Fig. S1)**.

### Single Molecular Fluorescent In-Situ Hybridization (smFISH)

To define mRNA expression patterns of *ACE2* and *TMPRSS2* in human pancreata, smFISH was performed using the RNAscope® Multiplex Fluorescent V2 kit (Advanced Cell Diagnostics, Newark, CA) in FFPE tissue cross-sections (5μm) from six non-diabetic, SARS-CoV-2 negative human organ donors from nPOD (**Table S2, Fig. 1E-G**, and **Fig. S2A**). Slides were baked at 60°C for 1 hour, followed by dehydration with xylene for 5 minutes x 2 and 100% ethanol for 2 minutes at room temperature (RT). Next, slides were air dried at 60°C for 5 minutes, treated with hydrogen peroxide for 10 minutes at RT, and washed 4 times with ddH_2_O, followed by antigen-retrieval at 99°C for 15 minutes. After another wash with ddH_2_O at RT for 15 seconds, slides were incubated at 100% ethanol for 3 minutes, air dried and then treated with protease plus for 30 minutes at 40°C. Next, slides were hybridized with probes for *ACE2* (Advanced Cell Diagnostics) and *TMPRSS2* (Advanced Cell Diagnostics) and detected using secondary TSA plus fluorophores (1:1500 dilution) according to the manufacture’s protocol (Perkin Elmer, Waltham, MA). Slides were immediately washed with 1X PBS and PBS containing 2% FBS for 5 minutes, blocked with donkey serum for 30 minutes, and incubated with ready to use (RTU) guinea pig polyclonal anti-insulin (no dilution; Agilent Santa Clara, CA) and/or mouse monoclonal anti-CD34 antibody (1:1,000 dilution, Novus Biologicals) overnight. The following morning, slides were washed with 1X PBS and PBS containing 2% FBS for 5 minutes and probed using either Alexa Fluor (AF)-488 goat anti-guinea pig IgG (1:500 dilution, Invitrogen, Carlsbad, CA) or AF-488 donkey anti-mouse IgG (1:1,000 dilution, Invitrogen) secondary antibodies. Finally, the slides were washed with PBS, counterstained with DAPI, and mounted with a coverslip using ProLong™ Gold antifade mounting media (Thermo Fisher, Rockford, IL). Images were acquired using an LSM800 confocal microscope (Carl Zeiss, Germany).

### Tissue Homogenization

Pancreas tissues were homogenized in modified radioimmunoprecipitation (RIPA) buffer (50 mM Tris-HCl pH 7.4, 150 mM NaCl, 5 mM EDTA, 1 mM EGTA, 1% NP-40, 0.5% sodium deoxycholate and 0.1% SDS) with freshly prepared protease and phosphatase inhibitor cocktails (Thermo Fisher Scientific) using a Tissue Tearor (BioSpec Inc., Bartlesville, OK). After homogenization, tissue was further disrupted with a Vibra-Cell™ sonicator (Sonics & Materials Inc., Newtown, CT) for 15 seconds, placed on ice for 15 minutes, then sonicated again and placed on ice for another 15 minutes. The tissue homogenates were centrifuged at 14,000 x *g* for 20 minutes at 4°C, and the supernatants were assayed for total protein concentration using a Pierce™ BCA Assay kit (Thermo Fisher Scientific) and stored at -80°C until use.

### Western Blotting

Fifty micrograms of protein lysates in Laemmli’s buffer containing 2.5% β-mercaptoethanol were boiled at 100°C for 5 minutes prior to loading into a 4-20% gradient Mini-PROTEAN TGX™ Stain-Free gel (Bio-Rad, Hercules, CA). After protein separation, gels were activated using a Gel DOC™ EZ imager (Bio-Rad), then transferred onto nitrocellulose membranes (LI-COR Biosciences, Lincoln, NE). Following protein transfer, membranes were scanned with the Gel DOC™ EZ imager, and total protein staining was visualized and quantified using Image Lab software version 5.2.1 (Bio-Rad). Then, membranes were washed and blocked for 1 hour at RT with Intercept™ Blocking Buffer (LI-COR Biosciences). Thereafter, the membranes were incubated at 4°C overnight with one of four primary antibodies (rabbit monoclonal anti-ACE2 (1:1,000 dilution, Abcam), rabbit polyclonal anti-ACE2 (1:500 dilution, Abcam), mouse monoclonal anti-ACE2 (1:1,000 dilution, R&D Systems), goat polyclonal anti-ACE2 (1:500 dilution, R&D Systems)) and mouse monoclonal anti-β-actin (1:10,000 dilution; Sigma-Aldrich, St. Louis, MO) in Intercept™ Antibody Diluent (LI-COR Biosciences). The membranes were then washed with Tris-buffered saline containing 0.1% Tween 20 (TBST) three times at 5 minute intervals, incubated with secondary antibodies (IRDye 800CW goat anti-rabbit IgG (1:30,000 dilution), IRDye 800CW goat anti-mouse IgG (1:30,000 dilution), IRDye 800CW donkey anti-goat IgG (1:30,000 dilution), or IRDye 680LT donkey anti-mouse (1:40,000 dilution), all from LI-COR Biosciences) for 1 hour at RT. The membranes were washed three times with TBST at 5-minute intervals. Immunoreactive bands were visualized and densitometrically analyzed using Odyssey infrared scanner and Image Studio software version 3.1 (LI-COR Biosciences) (**Fig. 2B** and **Fig. S2B**).

### Immunohistochemistry

FFPE pancreas, duodenum, and kidney tissues were sectioned (4μm), deparaffinized, rehydrated by serially passing through changes of xylene and graded ethanol, subjected to heat induced antigen retrieval in 10mM Citra pH 6, and blocked with avidin, biotin, and goat serum. For single-stained kidney, duodenum, and pancreas tissue sections (**Fig. 2C** and **Fig. S2D**) and for SARS-CoV-2 NP single stained lung and pancreas sections (**Fig. 4E-F** and **Fig. S4**), slides were incubated overnight at 4°C with one of four primary antibodies against ACE2 (rabbit monoclonal anti-ACE2, 1:200 dilution (Abcam); rabbit polyclonal anti-ACE2, 1:2,000 dilution (Abcam); mouse monoclonal anti-human ACE2, 1:100 dilution, (R&D Systems); goat polyclonal anti-ACE2, 1:100 dilution, (R&D Systems)) or a primary antibody against SARS-CoV-2 NP (mouse monoclonal anti-SARS-CoV-2 NP, 1:50 dilution, Invitrogen). Slides were washed, then incubated for 30 minutes at RT with biotinylated secondary antibodies (biotinylated goat anti-rabbit IgG, 1:200 dilution (Vector Laboratories, Burlingame, CA); biotinylated horse anti-mouse IgG, 1:200 dilution (Vector Laboratories); biotinylated rabbit anti-goat, 1:200 dilution (Vector Laboratories)), and developed with ImmpactDAB (Vector Laboratories) followed by hematoxylin counterstain.

For peptide blocking experiments (**Fig. S2E**) the primary antibody (monoclonal rabbit anti-ACE2, 1:100 dilution (Abcam, Cambridge, MA)) was incubated with 1mg/mL ACE2 peptide (Abcam) for one hour at RT, before applying to pancreas slides for overnight incubation at 4°C. Thereafter, IHC methodology was carried out as described for single stained sections.

For double- and triple-stained slides, FFPE pancreas slides were prepared for heat induced antigen retrieval in Borg Decloaker RTU (BioCare Medical, Pacheco, CA) followed by 3% H_2_O_2_. After washing, tissues were blocked with Background Sniper (BioCare Medical) followed by staining. For insulin and ACE2 double-staining (**Fig. 3A** and **Fig. S3A**), blocked slides were incubated for 20 minutes at RT with the first primary antibody (rabbit monoclonal anti-ACE2, 1:200 dilution (Abcam)), then washed and incubated with MACH 2 Double Stain Kit 1 (BioCare Medical) for 20 minutes at RT. Slides were washed and developed using DAB Chromogen solution (BioCare Medical), then subjected to a second round of heat-induced antigen retrieval in Borg Decloaker RTU, followed by 3% H_2_O_2_. After washing, slides were again blocked with Background Sniper, washed, and incubated with the second primary antibody (rabbit monoclonal anti-insulin, 1:2,000 dilution (Abcam)) for 30 minutes at RT. After washing, slides were incubated with MACH 2 Double Stain Kit 2 for 30 minutes at RT, washed, and developed with Warp Red Chromogen solution (BioCare Medical) followed by Hematoxylin counterstain. For ACE2, insulin and glucagon triple-staining (**Fig. 4D**), blocked slides were incubated for 20 minutes at RT with a primary antibody cocktail (mouse monoclonal anti-glucagon, 1:1,000 dilution (Abcam) plus rabbit monoclonal anti-ACE2, 1:200 dilution (Abcam)), then washed and incubated with MACH 2 Double Stain Kit 1 (BioCare Medical) for 20 minutes at RT. Slides were washed and developed using DAB Chromogen solution for ACE2 visualization followed by Ferangi Blue Chromogen solution (BioCare Medical) for glucagon visualization. Slides were then subjected to a second round of heat-induced antigen retrieval in Borg Decloaker RTU, followed by 3% H_2_O_2_. After washing, slides were again blocked with Background Sniper, washed, and incubated with the third primary antibody (rabbit monoclonal anti-insulin, 1:2,000 dilution (Abcam)) for 30 minutes at RT. After washing, slides were incubated with MACH 2 Double Stain Kit 2 for 30 minutes at RT, washed, and developed with Warp Red Chromogen solution (BioCare Medical) followed by Hematoxylin counterstain.

Following single-, double-, or triple-IHC staining, whole slides were scanned at an absolute magnification of 20x using an Aperio CS2 Scanscope (Leica/Aperio, Vista, CA), and stored in the nPOD online digital pathology database (eSLIDE version 12.4.0.5043, Leica/Aperio).

### Immunofluorescence

For immunofluorescence staining, FFPE pancreas sections were sectioned (4μm), deparaffinized, and rehydrated with antigen retrieval in 10mM Citra pH 6 and blocking as described above for IHC. Slides were incubated overnight at 4°C with primary antibodies: a) monoclonal rabbit anti-ACE2 (1:100 dilution; Abcam) (**Fig. 3C-E**), monoclonal rabbit anti-ACE2 (1:100 dilution; Abcam), polyclonal guinea pig anti-insulin RTU antibody (undiluted; Agilent), and monoclonal mouse anti-glucagon (dilution; 1:20,000 Abcam) (**Fig. 3F**) or b) monoclonal rabbit anti-ACE2 (1:100 dilution; Abcam) and monoclonal mouse anti-CD34 (dilution 1:1,000; Novus Biologicals, Centennial, CO) (**Fig. S3D**). Slides were washed, then incubated for 45 minutes at RT in the dark with secondary antibodies: a) goat anti-rabbit IgG-AF555, goat anti-mouse IgG-AF488, and goat anti-guinea pig IgG-AF647, or b) goat anti-rabbit IgG-AF594 and goat anti-mouse IgG-AF488 (all from Invitrogen). Slides were washed, then counterstained with DAPI and viewed using a Keyence BZ-X700 automated fluorescence microscope.

### H&E Staining

H&E staining was performed on FFPE pancreas tissues sections (4μm) from the three COVID-19 autopsy subjects according to standard methodology. Whole slides were scanned using an Aperio CS2 Scanscope (Leica/Aperio, Vista, CA), and stored in the nPOD online digital pathology database (eSLIDE version 12.4.0.5043, Leica/Aperio).

## QUANTIFICATION AND STATISTICAL ANALYSIS

For scRNAseq analysis, n represents the number of cells as indicated in the figure legends. Analyses were performed in R as described in METHOD DETAIL above. Differences in the average gene expression levels between pancreatic cell subsets or within each cell subset from non-diabetic donors versus donors with T2D were compared using Wilcoxon rank sum tests, requiring a minimum 1.19-fold change between the two groups and expression in at least 10% of cells from either group. Bonferroni corrections were used to adjust for multiple comparisons. Paired t-tests were used to compare proportions of cells with detectable gene expression. *P* values < 0.05 were considered significant. Violin plot limits show maxima and minima, and the dots represent individual data points. smFISH data was not quantitatively evaluated. For the quantification of ACE2 protein expression throughout the human lifespan, digitized images of ACE2 and insulin co-stained slides were analyzed using the HALO quantitative image analysis platform V3.0.311.262 (Indica Labs, Inc, Corrales, NM) (**Fig. 3A**). The annotation pen tool was used to outline the tissue section to determine total tissue area (mm^2^). The Area quantification algorithm v2.1.3 based on red, blue, green (RBG) spectra was employed to detect ACE2 positive tissue area stained for 3,3’-Diaminobenzidine (DAB, brown). The algorithm detected DAB IHC positivity and calculated percentage of ACE2 positive area per total tissue area. Donors (n=36) were binned into six age groups as described in the Results section. Data were analyzed in GraphPad Prism v8.3 (GraphPad Software, San Diego, CA) by one-way ANOVA followed by Tukey’s post hoc test for multiple comparisons with significance defined as P < 0.05 and graphed with the median percent ACE2 area shown for each group in a box and whisker plot. The remaining IF and IHC data were not quantitatively evaluated.

## KEY RESOURCES TABLE

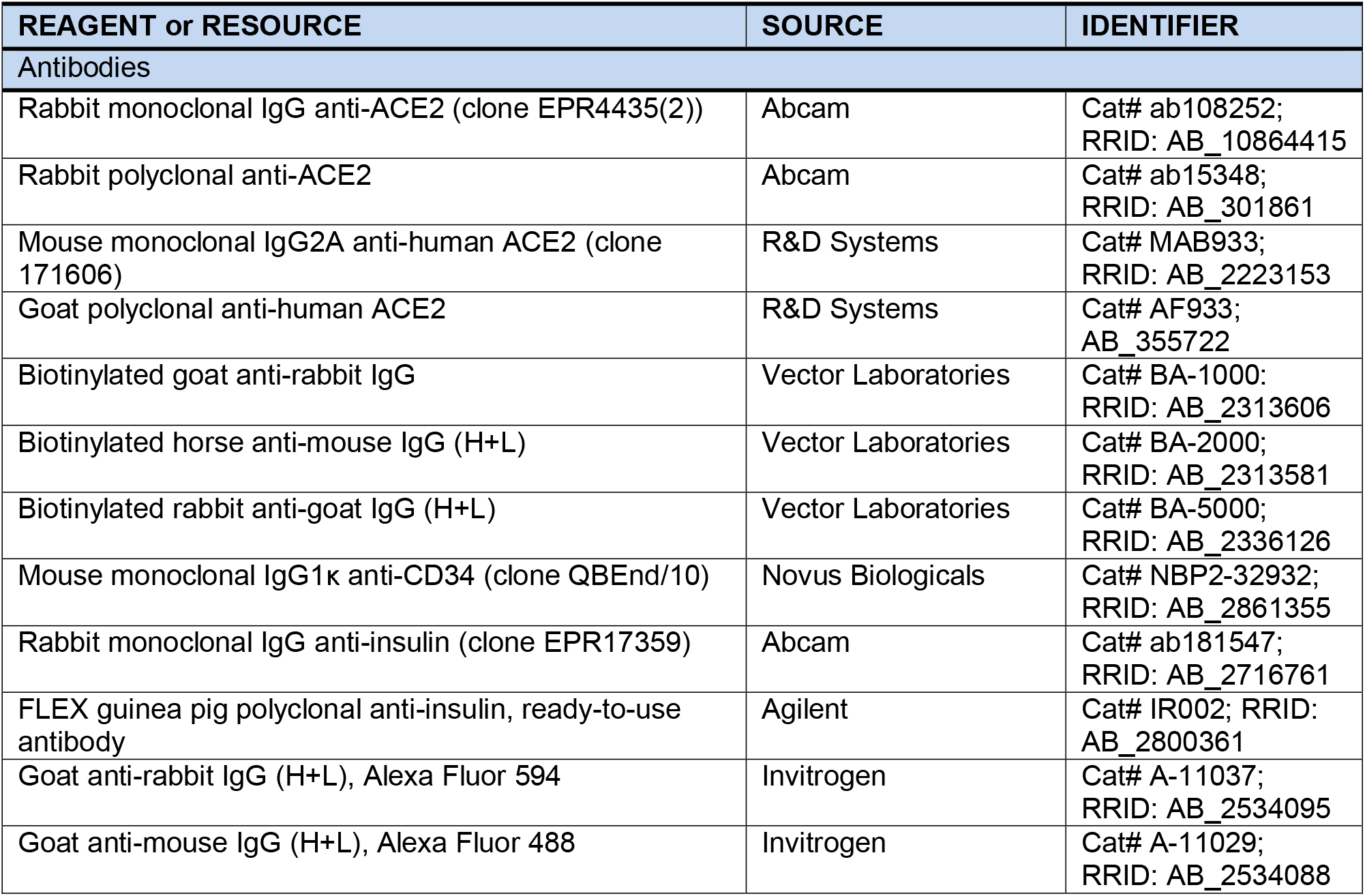

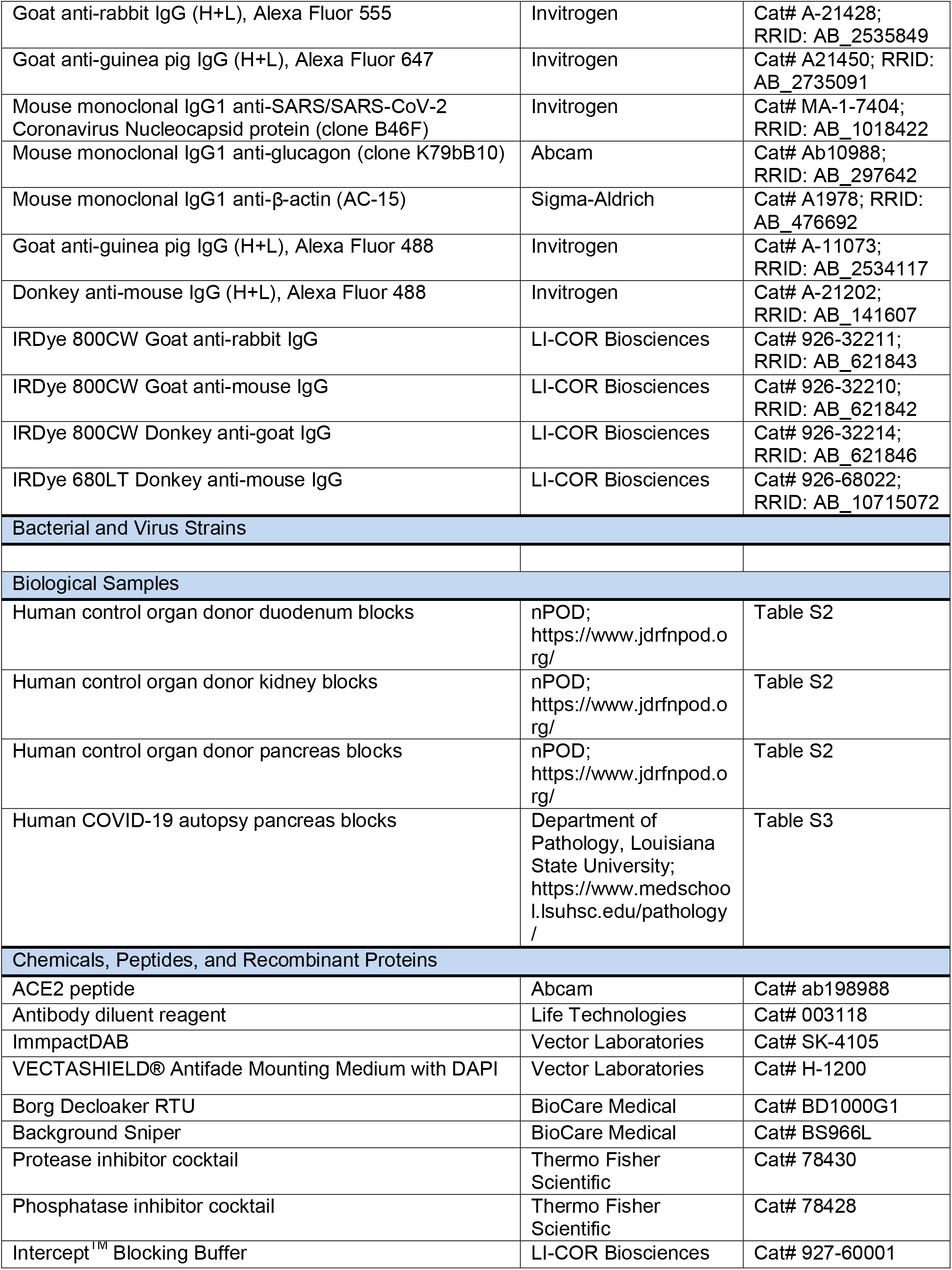

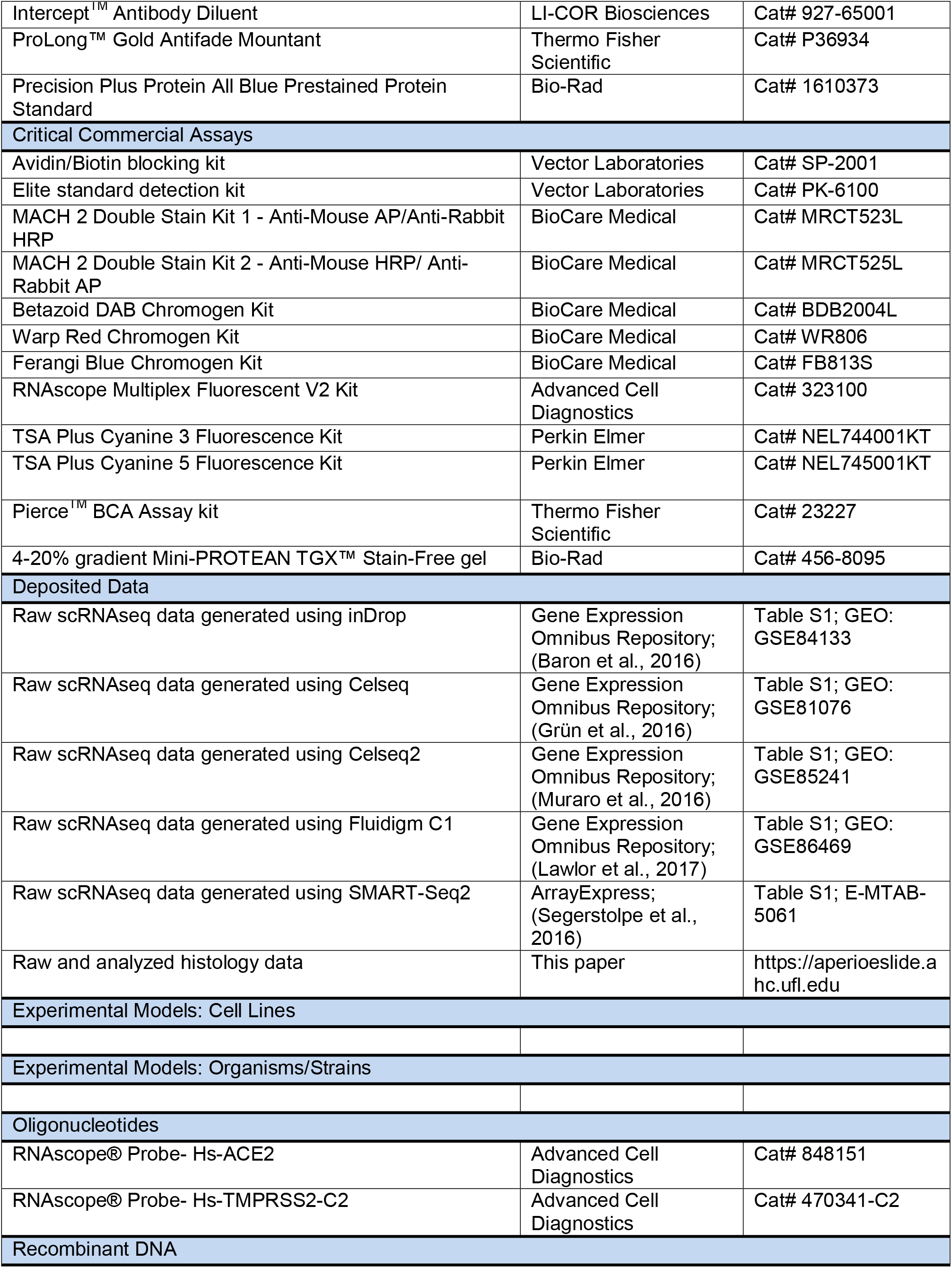

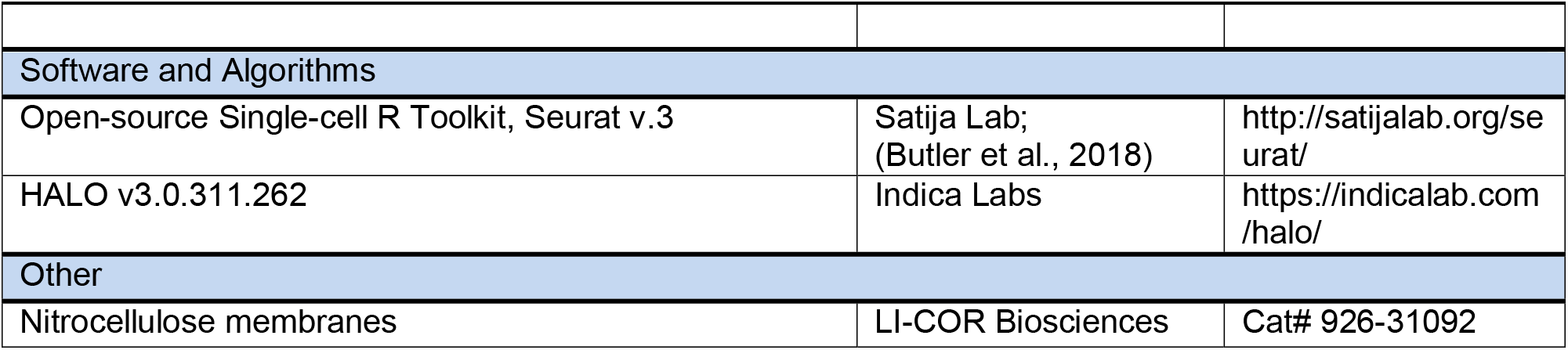

