## Supplemental Materials for "ACE2 and SARS-CoV-2 Expression in the Normal and COVID-19 Pancreas"

**Supplemental Tables**

**Table S1. Gene expression by donor group and pancreatic cell type.** From an integrated analysis of five public scRNAseq datasets (GSE84133 (Baron et al., 2016), GSE81076 (Grün et al., 2016), GSE85241 (Muraro et al., 2016), GSE86469 (Lawlor et al., 2017), and E-MTAB-5061 (Segerstolpe et al., 2016)), the number and percentage (presented as n (%)) of cells expressing *ACE2*, *TMPRSS2*, *TMPRSS4*, *CTSL*, *ADAM17*, and *TMPRSS11D* are listed for each cell type from isolated islets from control (Cont) versus type 2 diabetes (T2D) donors.

See also **Figure 1A-D** and **Figure S1**.

| Gene | Donor Group | Cell Type | | | | | | | | | | | | |
|---|---|---|---|---|---|---|---|---|---|---|---|---|---|---|
| | | Alpha | Beta | Delta | Epsilon | Gamma | Acinar | Ductal | Endothelial | Activated Stellate | Quiescent Stellate | Schwann | Macrophage | Mast |
| ***ACE2*** | Cont | 15 (0.40) | 17 (0.57) | 7 (0.83) | 0 (0.00) | 6 (1.41) | 70 (4.11) | 81 (5.54) | 0 (0.00) | 1 (0.26) | 2 (1.17) | 0 (0.00) | 1 (1.54) | 0 (0.00) |
| | T2D | 1 (0.12) | 2 (0.29) | 1 (0.57) | 0 (0.00) | 3 (1.52) | 13 (8.07) | 40 (8.13) | 0 (0.00) | 1 (1.19) | 0 (0.00) | 0 (0.00) | 0 (0.00) | 0 (0.00) |
| ***TMPRSS2*** | Cont | 624 (16.55) | 163 (5.46) | 24 (2.86) | 1 (3.85) | 18 (4.22) | 915 (53.73) | 739 (50.55) | 8 (2.88) | 10 (2.56) | 2 (1.17) | 3 (14.29) | 5 (7.69) | 2 (4.17) |
| | T2D | 271 (32.07) | 73 (10.52) | 9 (5.17) | 0 (0.00) | 11 (5.56) | 115 (71.43) | 289 (58.74) | 8 (44.44) | 8 (9.52) | 0 (0.00) | 2 (50.00) | 0 (0.00) | 0 (0.00) |
| ***TMPRSS4*** | Cont | 134 (3.55) | 151 (5.06) | 16 (1.91) | 1 (3.85) | 11 (2.58) | 31 (1.82) | 46 (3.15) | 6 (2.16) | 5 (1.28) | 1 (0.58) | 3 (14.29) | 1 (1.54) | 3 (6.25) |
| | T2D | 89 (10.53) | 81 (11.67) | 9 (5.17) | 0 (0.00) | 5 (2.53) | 13 (8.07) | 40 (8.13) | 7 (38.89) | 7 (8.33) | 0 (0.00) | 1 (25.00) | 0 (0.00) | 0 (0.00) |
| ***CTSL*** | Cont | 1375 (36.47) | 1006 (33.70) | 206 (24.55) | 11 (42.31) | 145 (33.96) | 353 (20.73) | 344 (23.53) | 98 (35.25) | 228 (58.46) | 123 (71.93) | 9 (42.86) | 42 (64.62) | 7 (14.58) |
| | T2D | 617 (73.02) | 409 (58.93) | 97 (55.75) | 3 (75.00) | 147 (74.24) | 91 (56.52) | 318 (64.63) | 9 (50.00) | 70 (83.33) | 7 (77.78) | 3 (75.00) | 13 (92.86) | 2 (25.00) |
| ***ADAM17*** | Cont | 709 (18.81) | 512 (17.15) | 140 (16.69) | 3 (11.54) | 97 (22.72) | 409 (24.02) | 483 (33.04) | 80 (28.78) | 116 (29.74) | 12 (7.02) | 8 (38.10) | 24 (36.92) | 6 (12.50) |
| | T2D | 370 (43.79) | 214 (30.84) | 61 (35.06) | 2 (50.00) | 97 (48.99) | 90 (55.90) | 316 (64.23) | 12 (66.67) | 39 (46.43) | 3 (33.33) | 2 (50.00) | 6 (42.86) | 3 (37.50) |
| ***TMPRSS11D*** | Cont | 17 (0.45) | 2 (0.07) | 3 (0.36) | 0 (0.00) | 0 (0.00) | 4 (0.23) | 5 (0.34) | 0 (0.00) | 0 (0.00) | 0 (0.00) | 1 (4.76) | 0 (0.00) | 0 (0.00) |
| | T2D | 16 (1.89) | 7 (1.01) | 0 (0.00) | 0 (0.00) | 5 (2.53) | 2 (1.24) | 8 (1.63) | 0 (0.00) | 1 (1.19) | 0 (0.00) | 0 (0.00) | 0 (0.00) | 0 (0.00) |

**Table S2. Demographic information, pancreatic histopathology report, and assay(s) performed for each donor organ.** Tissues from 41 non-diabetic, SARS-CoV-2 negative donors were selected from the nPOD biobank and subjected to assays as detailed in the far right column. Abbreviations: nPOD ID, Network for Pancreatic Organ donors with Diabetes Identification Number; yrs, years; BMI, body mass index; IHC, chromogen-based immunohistochemistry; IF, immunofluorescence; smFISH, single molecule fluorescence *in situ* hybridization.

*Three donors had hypertension in their anamnesis, but none of the donors evaluated received ACE inhibitors or angiotensin II receptor blockers during their terminal hospitalization stay.

See also **Figure 1E-G**, **Figure 2B-C**, **Figure 3**, **Figure S2**, and **Figure S3**.

| nPOD ID | Age (yrs) | Sex | Race/Ethnicity | BMI (kg/m$^2$) | Histopathology by nPOD | Technique Performed |
|---|---|---|---|---|---|---|
| 6012* | 68 | Female | Caucasian | 23.7 | Ins+/Gluc+ normal islets, high density tail. Mild chronic pancreatitis, IPMN in PanHead (gastric type). Moderate exocrine fatty infiltrate. | IHC |
| 6013 | 65 | Male | Caucasian | 24.2 | Ins+/Gluc+ normal islets. Focal, moderate ductular metaplasia. | IHC, IF, smFISH |
| 6017* | 59 | Female | Caucasian | 24.8 | Ins+/Gluc+ normal islets. Fatty infiltration-mild | IHC |
| 6021* | 72 | Female | Hispanic | 24.5 | Ins+/Gluc+ normal islets. Extra-acinar islets. Low Ki67. Mucinous ductal dysplasia. Multi-focal mild acinar atrophy and fatty infiltration. | IHC, IF |
| 6057 | 22 | Male | Caucasian | 26 | Ins+/Gluc+ normal islets, High Ki67+ most compartments | IHC |
| 6092 | 0.5 | Female | African Am | 13.8 | Ins+ normal. High Ki67+ islets and acinar. | IHC |
| 6106 | 2.9 | Male | Caucasian | 17.4 | Ins+/Gluc+ normal islets. Low Ki67 acinar and islet but multifocal, mild duct proliferation. | IHC |
| 6125 | 0.42 | Male | Caucasian | 18.9 | Ins+/Gluc+ normal islets. High acinar Ki67+. | IHC |
| 6130 | 5.2 | Male | Caucasian | 18.5 | Ins+ normal islets; low Ki67 in acinar and islets. No infiltrates. | IHC |
| 6137 | 8.9 | Female | Hispanic | 24.2 | Ins+ islets. Occ. islet with 3 or more Ki67+ cell. Mild fatty infiltrate. No inflammatory infiltrates. | IHC |
| 6164 | 0 | Male | Caucasian | 16.5 | Ins+/Gluc+ islets, plentiful. Multifocal lymphoid (high CD3+) aggregates or early PLN. High Ki67 islets and acinar regions. | IHC |
| 6168 | 51 | Male | Hispanic | 25.2 | Ins+/Gluc+ islets, all sizes. Normal density. Low Ki67. Variable fatty infiltrates acinar regions. | IHC, IF |
| 6178 | 24.5 | Female | Caucasian | 27.5 | Ins+/Gluc+ normal islets. Low Ki67. No infiltrates. | IHC |
| 6187 | 0.4 | Male | Caucasian | 17.1 | Ins+/Gluc+ normal islets, numerous. No inflammation. Low islet Ki67, moderate acinar Ki67. | IHC, IF |
| 6218 | 0.08 | Female | African Am | 17.2 | Ins+/Gluc+ islets, normal. No abnormalities observed. | IHC, IF |
| 6222 | 0.17 | Male | Caucasian | 16.4 | Ins+/Gluc+ islets, numerous. High exocrine Ki67 with focal high duct Ki67+. Islets with Ki67+ cells in occ. islets head and body with more in tail region | IHC |
| 6229 | 31 | Female | Caucasian | 26.9 | Ins+/Gluc+ islets, no abnormalities observed. Occasional islet hyperemia. | IHC |

| | | | | | | |
|---|---|---|---|---|---|---|
| 6232 | 14 | Female | Caucasian | 20.8 | Ins+/Gluc+ islets, numerous. No significant findings. 0-4 Ki67+ cells per islet, some non-insulin, as expected for this age. Low acinar Ki67. | IHC |
| 6251 | 33 | Female | Caucasian | 29.5 | Ins+/Gluc+ islets, numerous, including single cells. Infrequent islet with 1-3 Ki67+ cells. No significant lesions. | IHC, IF, Western Blot |
| 6282 | 14 | Male | Caucasian | 41.9 | Ins+/Gluc+ islets, numerous, small to large islet size range. High islet Ki67+ (2-5+ cells/islet). | IHC, IF |
| 6290 | 58 | Male | Caucasian | 22.5 | Ins+/Gluc+ islets, numerous. Multifocal extra- and intracinar regions with moderate fat content. Very mild chronic pancreatitis, focal (PanTail). | IHC |
| 6313 | 0.25 | Male | Caucasian | 15.5 | Ins+/Gluc islets, mostly small, numerous islets. Moderate acinar and islet Ki67+. No CD3 infiltrates observed. | IHC, IF, smFISH |
| 6315 | 1.6 | Male | African Am | 14.7 | Ins+/Gluc+ islets, normal sizes and morphologies. Moderate acinar Ki67+ while islets have low Ki67+. No infiltrates or other significant lesions. | IHC |
| 6348 | 0.03 | Female | African Am | 11.9 | Ins+/Gluc+ islets, clusters, single cells. Well formed acini and lobules throughout. Very high Ki67 acini and islets. | IHC |
| 6353 | 13 | Male | African Am | 28.3 | Ins+/Gluc+ islets, range of normal sizes with rare islet >500um. Moderate islet Ki67+. Mild extra-pancreatic fat. Low acinar Ki67+ and CD3+ cell numbers. | IHC,IF |
| 6356 | 1.58 | Female | Caucasian | 17.1 | Ins+/Gluc+ islets, normal range of sizes and morphologies. Very mild focal islet hyperplasia ventral lobe. Acinar Ki67+ cell numbers mild with very Ki67+ islets for donor age. | IHC, IF, smFISH |
| 6370 | 0 | Male | Caucasian | 11.4 | Ins+/Gluc+ islets, expected numbers and morphologies including numerous single cells and clusters. High exocrine Ki67+ cell numbers but infrequent Ki67+ beta cells. Minimal exocrine CD3+ cell numbers. | IHC |
| 6375 | 28.7 | Male | Caucasian | 31.8 | Ins+/Gluc+ islets, normal numbers and sizes. Highly variable islet Ki67+ numbers with some having very high Ki67+. Low acinar Ki67+. Low with focally increased intralobular fat. Low exocrine infiltrates. | IHC |
| 6376 | 0.6 | Female | Caucasian | 19.4 | Ins+/Gluc+ islets, expected range of small to medium sized islets with regular morphology. Moderate Ki67+ acinar cells with focally high numbers in tail region. Low islet Ki67+ cells. Tail region has multifocal, very mild CD3+ infiltrates. | IHC |
| 6387 | 15.6 | Male | Caucasian | 18.1 | Ins+/Gluc+ islets, density, sizes, and morphologies within normal range. Low Ki67 except for few islets with mild increases, often non-beta cell. Low (normal) exocrine infiltrates. | IHC |
| 6406 | 6.9 | Male | Caucasian | 16.8 | Ins+/Gluc+ islets within normal range of sizes, shapes and density per region. | IHC, IF, smFISH |
| 6415 | 10.9 | Male | African Am | 14.8 | Ins+/Gluc+ islets range of normal sizes, numbers and morphologies. Acinar and islet Ki67+ cell numbers low (0-2) to occasional low-moderate (3-10). | IHC |
| 6461 | 14.29 | Male | Caucasian | 18.5 | Ins+/Gluc+ islets, mostly small to medium with regular morphology. Rare possible Ins- islet and small islets or clusters of alpha cells. No significant lesions observed. | IF, smFISH |

| | | | | | | |
|---|---|---|---|---|---|---|
| 6467 | 13.83 | Male | Caucasian | 19.6 | Ins+/Gluc+ islets expected range of sizes, morphologies and densities with single beta-cells widely scattered. | IHC |
| 6479 | 21.67 | Female | Hispanic | 20.9 | Ins+/Gluc+ islets, normal range of sizes, morphologies and densities. Mild (head, body) to moderate lobular fatty infiltration. | IHC |
| 6488 | 4.6 | Female | Caucasian | 16.8 | Ins+/Gluc+ islets, normal range of sizes, morphologies and densities. Abundant single beta- and alpha-cells. Mild increase exocrine Ki67+ cell numbers due to mild leukocytic infiltrate as often seen. | IHC |
| 6493 | 18.84 | Male | Caucasian | 17.7 | Ins+/Gluc+ islets, normal sizes and morphologies including single endocrine cells in acinar regions. No significant lesions observed. | IHC, IF, Western Blot |
| 6495 | 9.6 | Male | Caucasian | 10.7 | Ins+/Gluc+ islets, normal range of sizes including some large (800um) with regular morphologies and densities per region. No significant lesions. | Western Blot |
| 6500 | 14.14 | Female | African Am | 19.7 | Ins+/Gluc+ islets, wide range of sizes and mostly spherical morphologies. No significant lesions. | IHC |
| 6516 | 20.75 | Male | Caucasian | 28.8 | Ins+/Gluc+ islets, normal range of sizes, numbers and shapes. No significant findings. | IF, smFISH |
| 6518 | 21.86 | Male | Caucasian | 23.8 | (Preliminary): islets plentiful, normal numbers, morphology. | IHC |

**Table S3. Demographic and clinical information for COVID-19 patients.** Pancreas was obtained at autopsy from three patients with fatal coronavirus disease 2019 (COVID-19). Abbreviations: yr, years; African Am, African American; BMI, body mass index; Type 2 DM, Type 2 diabetes; appx, approximately.

See also **Figure 4** and **Figure S4C**.

| | **COVID-19 Patient 1** | **COVID-19 Patient 2** | **COVID-19 Patient 3** |
|---|---|---|---|
| Age at Death (yr) | 72 | 45 | 71 |
| Sex | Male | Male | Male |
| Race | Caucasian | African Am | African Am |
| BMI ($kg/m^2$) | 14.9 | 49.0 | 26.3 |
| Diabetes Status | No diabetes | Type 2 DM | Type 2 DM |
| Diabetes Duration | ---- | unknown | unknown |
| Blood Glucose (mg/dL)<br>at admission<br>at death | <br>94<br>121 | <br>156<br>238 | <br>264<br>134 |
| Reported Days Ill Prior to Hospital Admission | Unknown (appx 1 week nursing home ) | 2 | 4 |
| Days from Admission to Death | 51 | 8 | 39 |

### Supplemental Figures

**Supplemental Figure 1. Expression patterns of SARS-CoV-2 associated genes. (A-B)** Violin plot showing *ACE2* and *TMPRSS2* normalized gene expression in β-cells from donors with (n=694 cells) and without type 2 diabetes (T2D, n=2,985 cells). *ACE2* and *TMPRSS2* expression were not significantly different between groups, Wilcoxon rank sum tests. **(C)** Violin plot showing the distribution of *TMPRSS4* normalized expression in non-diabetic human pancreas cells (n=12,185 cells). **(D)** Bar graphs showing the percentage of pancreatic cells with detectable *TMPRSS4* in donors with (n=2,705 cells) and without T2D (n=12,185 cells). **(E-G)** Violin plots showing the distribution of *TMPRSS11D*, *CTSL*, and *ADAM17* normalized expression in non-diabetic human pancreas cells (n=12,185 cells). **(H)** Violin plot showing *CTSL* normalized gene expression in β-cells from donors with (n=694 cells) and without type 2 diabetes (n=2,985 cells). *CTSL* expression was higher in beta cells from donors with T2D, Wilcoxon rank sum tests, adjusted $P = 8.94 \times 10^{-32}$, $***$ $p < 0.001$. Violin plot limits show maxima and minima, and the dots represent individual data points. See also **Table S1** and **Figure 1A-D**.

Supplemental Figure 1

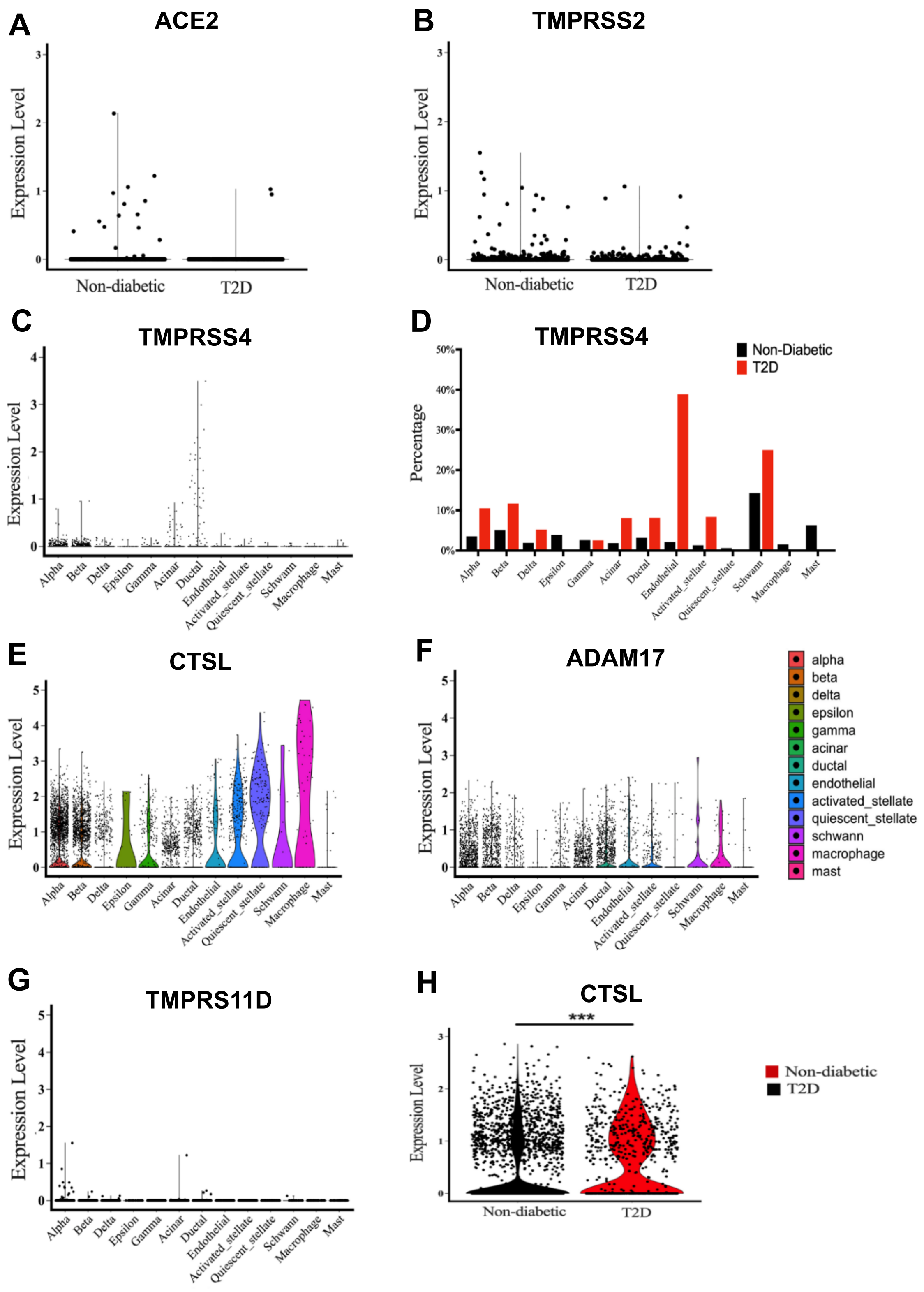

**Supplemental Figure 2. Detection of ACE2 mRNA and protein in human pancreatic tissue.** **(A)** Validation of smFISH probes in control human tissues. Single molecular fluorescent in-situ hybridization (smFISH) images of human duodenum, ileum, and kidney used as a positive controls to test the specificity of the probe sets used to determine the expression of *ACE2* and *TMPRSS2*. **(B)** Detection ACE2 in protein lysate from three organ donors. Full immunoblot images of data presented in Figure 2B. Membranes were incubated with primary antibodies against ACE2 (as indicated on the figure), washed, and incubated with the corresponded secondary antibody. Protein bands were visualized using the Odyssey Li-Cor infrared imager. The molecular weight ladder (Bio-Rad, Cat.No. 1610373) was run in each gel to confirm the correct size of the ACE2 band. (**C**) Total protein on the membrane for the corresponding blots shown above. Pancreas lysates were run in TGX stain-free gels (Bio-Rad). After electrophoresis, gels were activated under UV light for 5 minutes using a Gel Doc EZ imaging system (Bio-Rad) to label all tryptophan residues on proteins. Once photoactivated, proteins were transferred to nitrocellulose membranes. The stain-free signal from proteins on the membrane was imaged using the Gel Doc EZ system. **(D)** Validation of ACE2 antibodies for immunohistochemistry. IHC images of human duodenum and kidney were used as positive controls to test the specificity of four commercially available ACE2 antibodies as indicated on the figure. **(E)** ACE2 blocking peptide (Abcam, Cat# 198988) was used to confirm the specificity of Abcam rabbit monoclonal ACE2 antibody (Cat# ab108252). The image of pancreas tissue section stained for ACE2 showing absence of staining with the neutralized antibody. Scale bars: 500μm (D) and 300μm (E). See also **Table S2**, **Figure 1E-G** and **Figure 2**.

Supplemental Figure 2

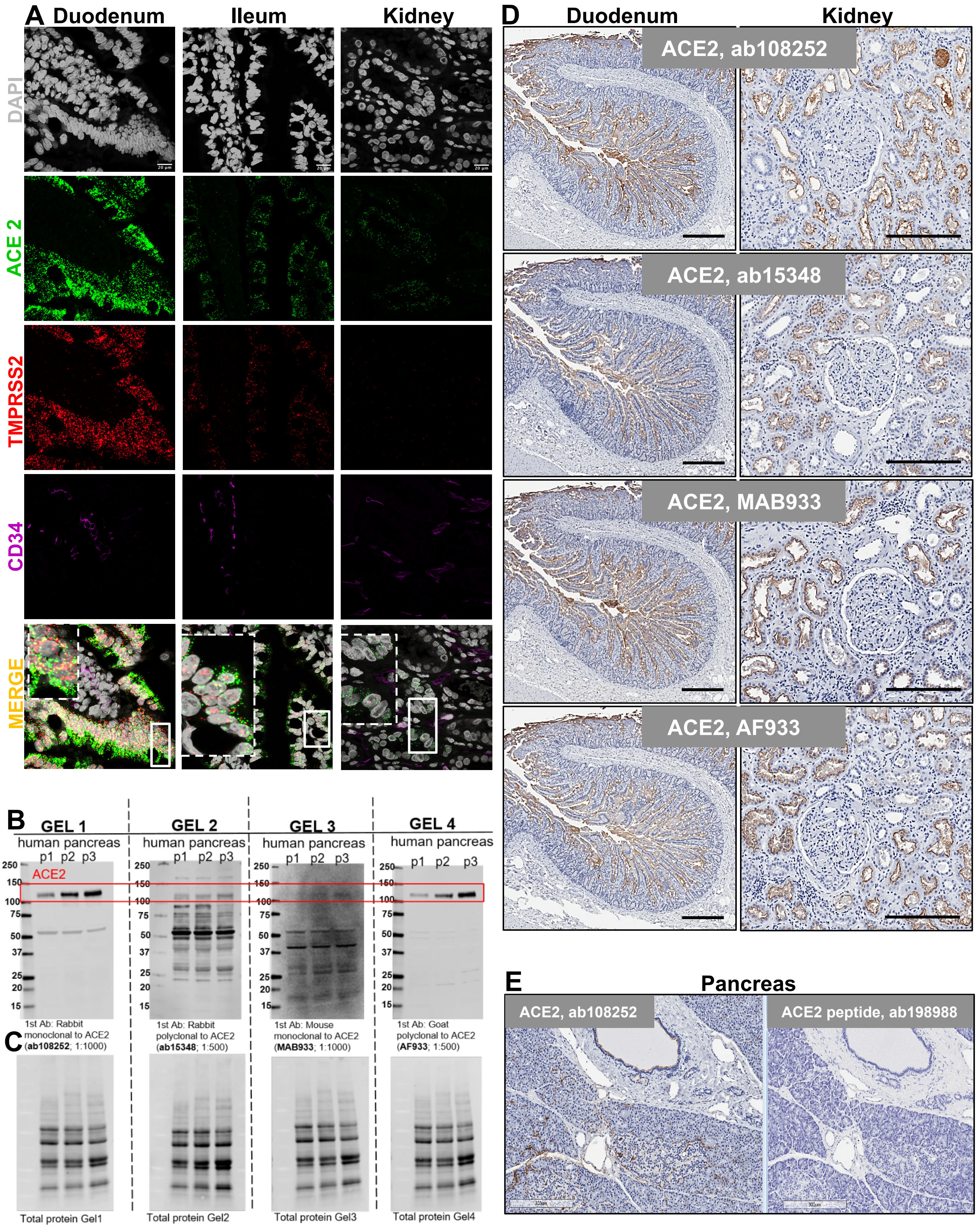

**Supplemental Figure 3. ACE2 is highly expressed in microvasculature of the human pancreas. (A**) Representative image of pancreatic tissue section from SARS-CoV-2 negative donor without diabetes stained for ACE2 and Insulin. ACE2 positivity was associated with microvasculature in the acinar **(B)** and islet regions **(C)** Scale bars: 200µm. **(D)** ACE2 expression by endothelial cells in the human pancreas. Representative immunofluorescence image of pancreatic tissue section stained for ACE2 (Abcam, ACE2 antibody Cat# ab108252) and endothelial cell marker CD34 (Novus Biological, mouse monoclonal CD34 antibody Cat# NBP2-32932). **(E-F)** Endothelial cells positive for both ACE2 and CD34 markers are shown. Scale bars: 50µm. See also **Table S2** and **Figure 3**.

Supplemental Figure 3

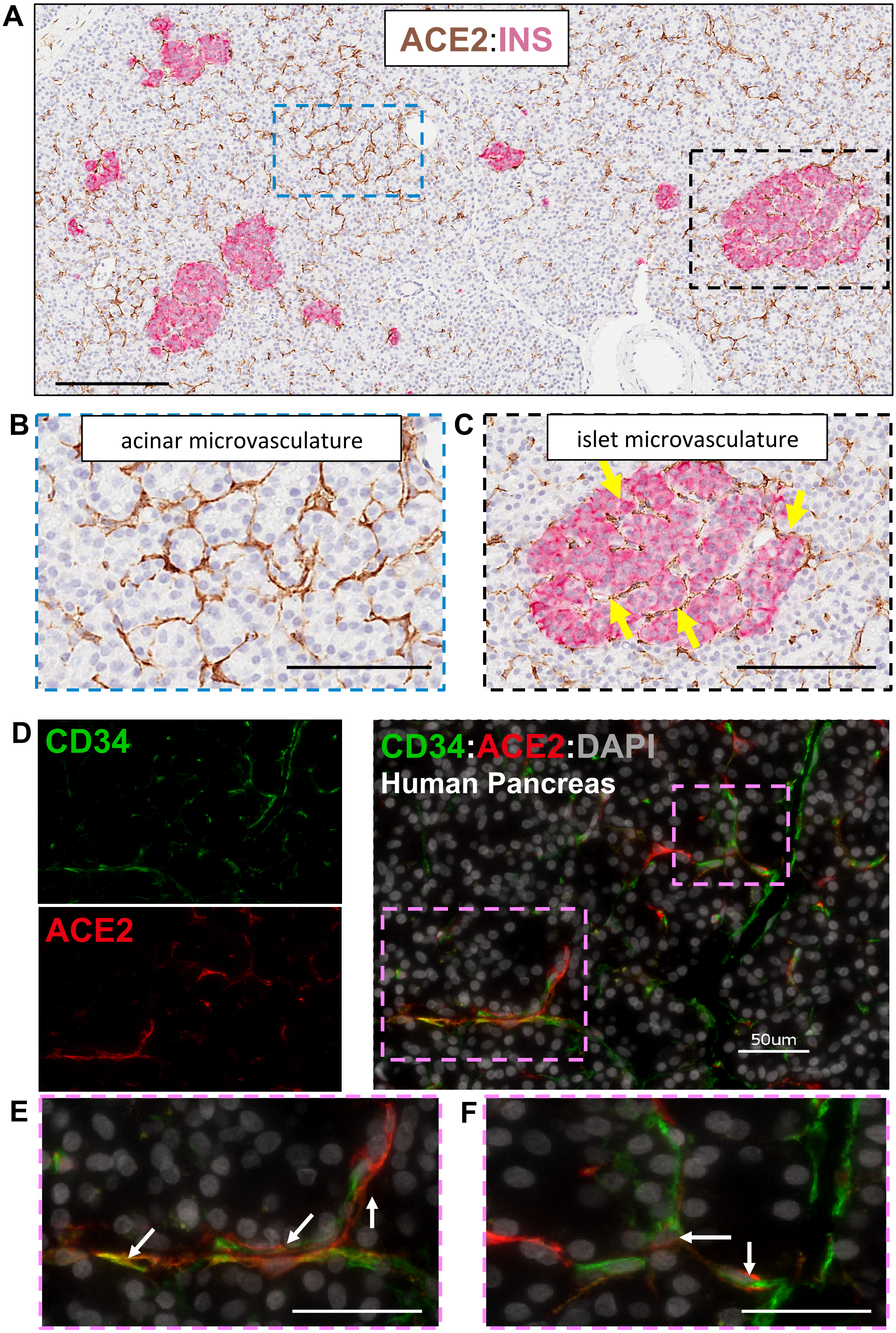

**Supplemental Figure 4. Detection of SARS-CoV-2 nucleocapsid protein (NP) in the pancreas of COVID-19 patient. (A-B)** IHC image of the human lung tissue sample known to be positive for SARS-CoV-2 used as positive control to test specificity of antibody against SARS-CoV-2 NP (clone B46F, Invitrogen, Cat# MA-1-7404). Scale bars: 50µm. **(C)** Tissue section from pancreas of COVID-19 patient 1 showing the presence of SARS-CoV-2 NP in ductal epithelial cells. Scale bar: 200µm. See also **Table S3** and **Figure 4**.

Supplemental Figure 4

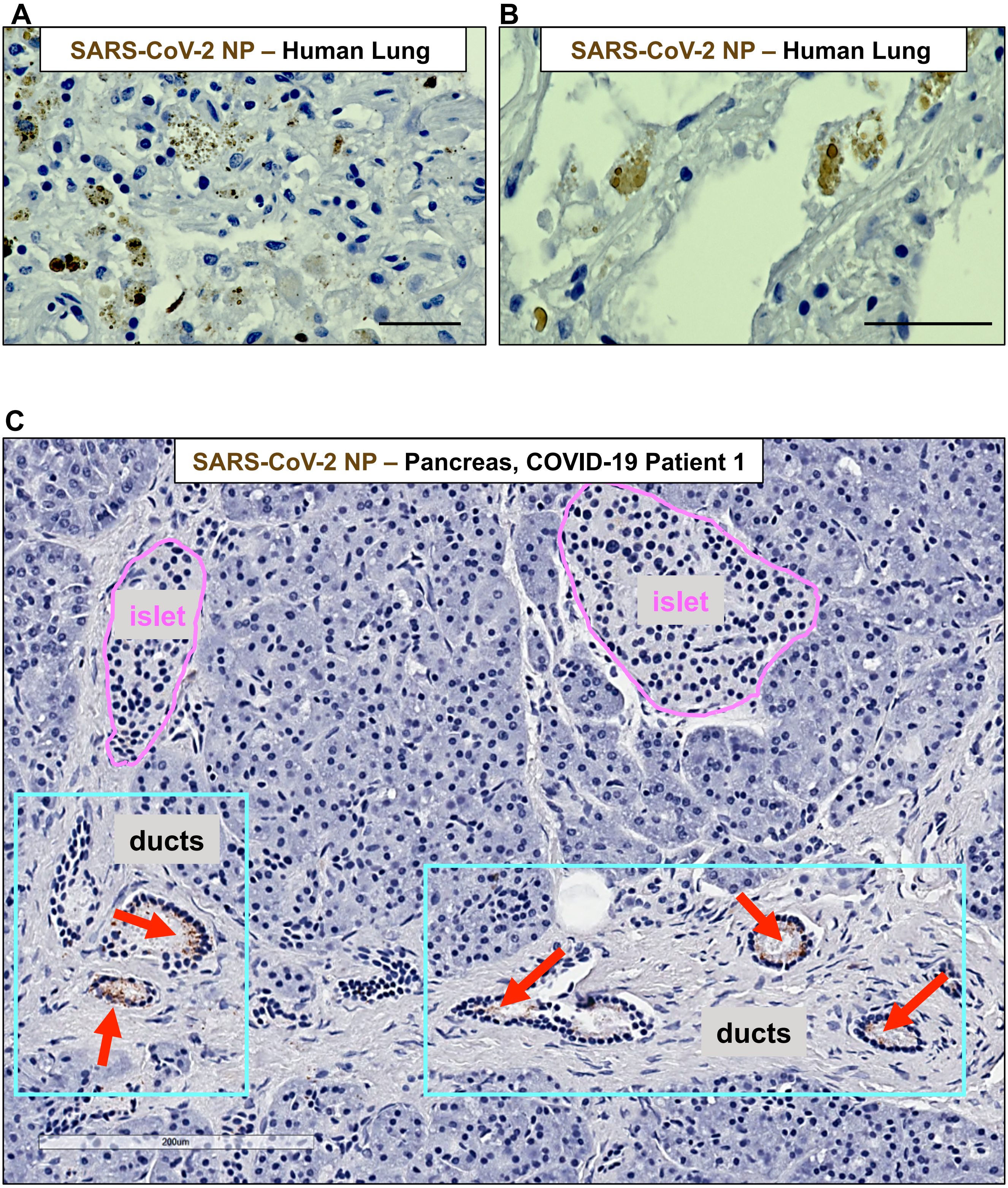
